# Development of intimin-enriched outer membrane vesicles (OMVs) as a vaccine to control intestinal carriage of Enterohemorrhagic *Escherichia coli*

**DOI:** 10.1101/2024.12.09.627534

**Authors:** Asja Garling, Cécile Goursat, Carine Seguy, Patricia Martin, Audrey Goman, Jean-Philippe Nougayrède, Éric Oswald, Frédéric Auvray, Priscilla Branchu

## Abstract

Enterohemorrhagic *Escherichia coli* (EHEC) are foodborne pathogens causing severe human infections including hemorrhagic colitis and hemolytic uremic syndrome (HUS), particularly in children. Ruminants are the main reservoir of EHEC which colonize their intestinal tract through a mechanism involving the bacterial outer membrane adhesin intimin. Vaccination of cattle has shown efficacy in reducing EHEC O157:H7 shedding in feces. However, most of these vaccines are based on purified proteins and/or require the addition of adjuvants, resulting in expensive vaccines that are not used by breeders. This study introduces the development of a new type of vaccine based on Outer Membrane Vesicles (OMVs) carrying the C-terminal domain of intimin (Int280). A vaccine which combines OMVs carrying luminal Int280 and OMVs displaying surface-exposed Int280 was produced using two addressing systems based on PelB peptide signal and Lpp-OmpA hybrid protein, respectively. This mixed vaccine was tested in a mouse model as a proof of concept using the murine host-specific intestinal pathogen *Citrobacter rodentium* which shares a similar intimin-based adhesion mechanism with EHEC. Vaccination of mice with OMV-Int280 elicited a strong anti-intimin IgG response. Interestingly, we observed a shortened *C. rodentium* fecal shedding duration in immunized mice compared to the control group. This OMVs-intimin vaccine therefore represents a promising candidate for the control of EHEC intestinal carriage and fecal shedding in ruminants.

**IMPORTANCE:** Enterohemorrhagic *Escherichia coli* (EHEC) are foodborne pathogenic bacteria causing intestinal infection that may lead to hemorrhagic colitis and hemolytic uremic syndrome (HUS) particularly in young children. There is no effective treatment, and antibiotics are contraindicated because they promote the development of HUS. Vaccination of ruminants, the main reservoir of EHEC, has been proposed as an important strategy to reduce the fecal shedding of EHEC to reduce transmission to humans. Outer Membrane Vesicles (OMVs) derived from *E. coli* are a highly attractive vaccine platform. Here, we produced OMVs enriched with the C-terminal part of the intimin (Int280). As a proof of concept, we used a mice model of *Citrobacter rodentium* colonization as a surrogate for EHEC intestinal colonization. Vaccination elicited antibodies against intimin and decreased the duration of fecal shedding of *C. rodentium*. Therefore, this OMV-Int280 vaccine is a promising candidate to control EHEC intestinal carriage and fecal shedding in ruminants.

## INTRODUCTION

Enterohemorrhagic *E. coli* (EHEC) represent a major public health issue. EHEC are foodborne human pathogens responsible for hemorrhagic colitis and hemolytic uremic syndrome (HUS) or even death (1, 2). HUS is the leading cause of acute renal failure, especially in children under the age of five (3). Since the 1990’s, EHEC have been regularly responsible for HUS outbreaks caused by the consumption of contaminated food (4–6). EHEC driven hemorrhagic colitis and HUS are triggered by the release of Shiga toxins (Stx) (1). Prior colonization of the intestinal tract by EHEC is required and generally involves the *Locus* of Enterocyte Effacement (LEE) which encodes the outer membrane (OM) adhesin named intimin (encoded by the *eae* gene) and its receptor Tir (7, 8). Intimin-Tir interaction mediates the intimate adhesion of EHEC to enterocytes and initiates a signaling cascade leading to the formation of actin- rich pedestals beneath adherent EHEC and the effacement of intestinal microvilli (8, 9). Such pedestals can be observed with the fluorescent actin staining (FAS) assay (10).

The main reservoir of EHEC is the digestive tract of ruminants (11, 12), where intimin is required, as in humans, for the intestinal colonization (13, 14). Colonized ruminants shed EHEC in their feces, which can contaminate foodstuffs, such as raw dairy products, ground meat or vegetables (1).

The sanitization treatments used by the food industry may not be sufficient to completely eliminate EHEC from food and prevent human infection, especially since the infectious dose is very low (< 100 colony forming units, CFU) (1, 15). Also, the contamination rate is usually very low, with a high heterogeneity within food products, rendering difficult the detection of EHEC. Furthermore, there is no specific treatment against EHEC infection in humans, and antibiotics are contraindicated due to their propensity to promote the production and release of Stx, thereby increasing the risk of developing HUS (16, 17). The occurrence of HUS cases and outbreaks has motivated the development of pre-harvest anti-EHEC strategies, notably the vaccination of ruminants (18, 19).

Extracellular vesicles, and particularly outer membrane vesicles (OMVs) derived from Gram-negative bacteria such as *E. coli*, have attracted increasing attention for vaccine development in the last decade due to their unique properties. OMVs are naturally produced, at low cost, by Gram-negative bacteria from the budding of their OM (20). Thus, they share similarities with the OM of the producing bacteria, in particular the presence of pathogen-associated molecular patterns (PAMPs) such as lipopolysaccharides (LPS) or outer membrane proteins (OMPs), which confer both immunogenic and auto-adjuvant properties (21). In addition, they cannot replicate, rendering them more attractive than attenuated or inactivated vaccines in terms of safety and reduced side effects (22). Finally, OMVs are a versatile tool that can be easily loaded with homologous and/or heterologous antigens on their surface or in their lumen (23–25).

The aim of this study was to develop a new generation of vaccine based on OMVs enriched with the carboxy-terminal 280 amino acids (AA) of intimin (Int280), that could be used in the future to inhibit fecal shedding of EHEC by ruminants. Ideally, vaccines should target both humoral and cellular immune responses to protect efficiently against pathogens (26). OMVs offer the possibility to elicit both responses. Indeed, an antigen presented at the surface of OMVs will be accessible to B cells to trigger mainly the humoral response (27, 28), while an antigen addressed in their lumen will induce mainly the cellular response (27, 29). To target EHEC, we developed a vaccine containing a mixture of OMVs with Int280 located either in their lumen or at their surface. To this aim, we constructed plasmids encoding Int280 recombinant protein fusions. For the production of luminal Int280-enriched OMVs, Int280 was fused to the signal sequence of the PelB protein. Its recognition by the general secretory (Sec) pathway is expected to promote its translocation into the *E. coli* periplasm, and thus into the lumen of OMVs (30, 31). For OMVs displaying Int280 at their surface, Int280 was fused to the hybrid Lpp-OmpA protein containing the signal sequence and the first 9 amino acids of the lipoprotein Lpp (AA1 to AA29) followed by OmpA transmembrane strands 3 to 7 (AA46 to AA159) (32).

As a proof of concept, we evaluated the efficacy of this Int280-enriched OMVs vaccine using the mouse *Citrobacter rodentium* infection model. Indeed, *C. rodentium* is a natural mouse pathogen that possesses an intimin-mediated adhesion mechanism similar to EHEC and thus reproduces ruminant intestinal colonization by EHEC (33–35).

## RESULTS

### Int280 is delivered into OMVs fractions by the PelB- and Lpp-OmpA-based systems

Two different systems based on the PelB signal sequence or Lpp-OmpA hybrid protein were used to address the 280 amino acids (AA) of the C-terminal domain of *C. rodentium* intimin (Int280) in the lumen or to the surface of OMVs, respectively (Fig. 1A).

**FIG 1.**
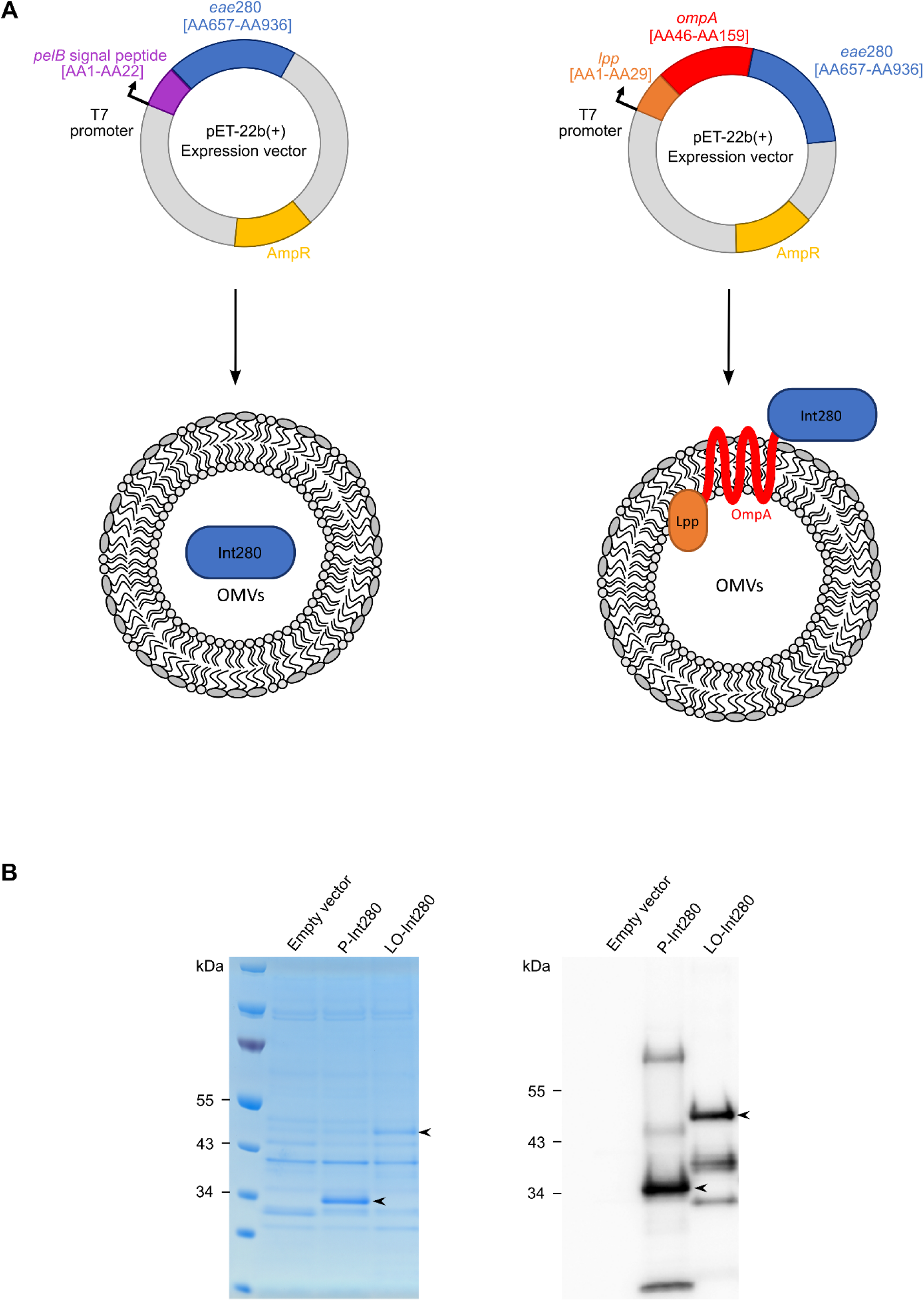
Addressing of *C. rodentium* Int280 to the lumen and surface of OMVs using PelB- and Lpp-OmpA-based systems. (**A**) Schematic representation of the recombinant plasmids and corresponding OMVs containing Int280 in their lumen or on their surface. The pET-22b(+) expression vector was used to express the PelB-Int280 (P-Int280) or Lpp-OmpA-Int280 (LO-Int280) fusion proteins under control of the T7 promoter. The PelB signal peptide addresses Int280 in the lumen of OMVs (left) while the hybrid protein Lpp-OmpA addresses Int280 to the surface of OMVs (right). The amino acids (AA) of each protein fragment incorporated in the fusion proteins are indicated. (**B**) Detection of P-Int280 and LO-Int280 in OMVs fractions. OMVs were obtained from *E. coli* BL21(DE3) Δ*ompF* containing plasmid pET-22b(+) (“Empty vector”), pAGA12 (“P-Int280”) or pAGA16 (“LO-Int280”). Bacterial cultures were induced with 1 mM of IPTG. An equivalent amount of OMVs (as determined by total protein assay) was loaded in each lane for analysis by SDS-PAGE stained with Coomassie Blue (10 µg OMVs extracts) (left panel) and Western blotting (2 µg OMVs extracts) (right panel). For Western blotting, membrane was probed with an anti-Int280 antibody. Locations of P-Int280 (30kDa) and LO-Int280 (43kDa) are indicated with arrowheads.

The resulting PelB-Int280 (P-Int280) and Lpp-OmpA-Int280 (LO-Int280) fusion proteins were expressed in the *E. coli* strain BL21(DE3) Δ*ompF* to avoid steric hindrance between the major porin OmpF (36) and Int280, and avoid seroconversion against OmpF. The P-Int280 and LO-Int280 fusion proteins were both detected in OMVs fractions, the former being more abundant than the latter (Fig. 1B).

### A functional Int280 is addressed by the Lpp-OmpA system to the OM surface of *E. coli*

The 3D structure of the antigen is crucial to induce an efficient immune response. To test if the Int280 fused to the hybrid Lpp-OmpA protein has a similar 3D structure to the full-length intimin, we tested its functionality by fluorescent actin staining (FAS) assay (10). For this test, we used the enteropathogenic *E. coli* (EPEC) O103:H2 strain E22 previously described to induce actin polymerization beneath the adherent bacteria (37). We constructed the plasmid pBRSK_LO-Int280 (pAGA43) where the expression of the *C. rodentium* Int280 fused to Lpp-OmpA is under the control of the native intimin gene promoter, allowing its production in the required conditions for the FAS assay. The mutant strain E22 Δ*eae* (which does not produce intimin) was transformed with either the plasmid pAGA43 or the empty vector pBRSK. The FAS phenotype was observed in cells infected with the E22 WT strain, but not in cells infected with the E22 Δ*eae* mutant strain hosting the empty vector (Fig. 2). In the latter case, the FAS phenotype was restored by complementing the E22 Δ*eae* mutant with plasmid pAGA43 expressing the LO-Int280 protein (Fig. 2). These results indicated that Int280, when fused to the hybrid Lpp-OmpA protein, was successfully exported to the bacterial surface of the EPEC E22 Δ*eae* strain and was functional.

**FIG 2.**
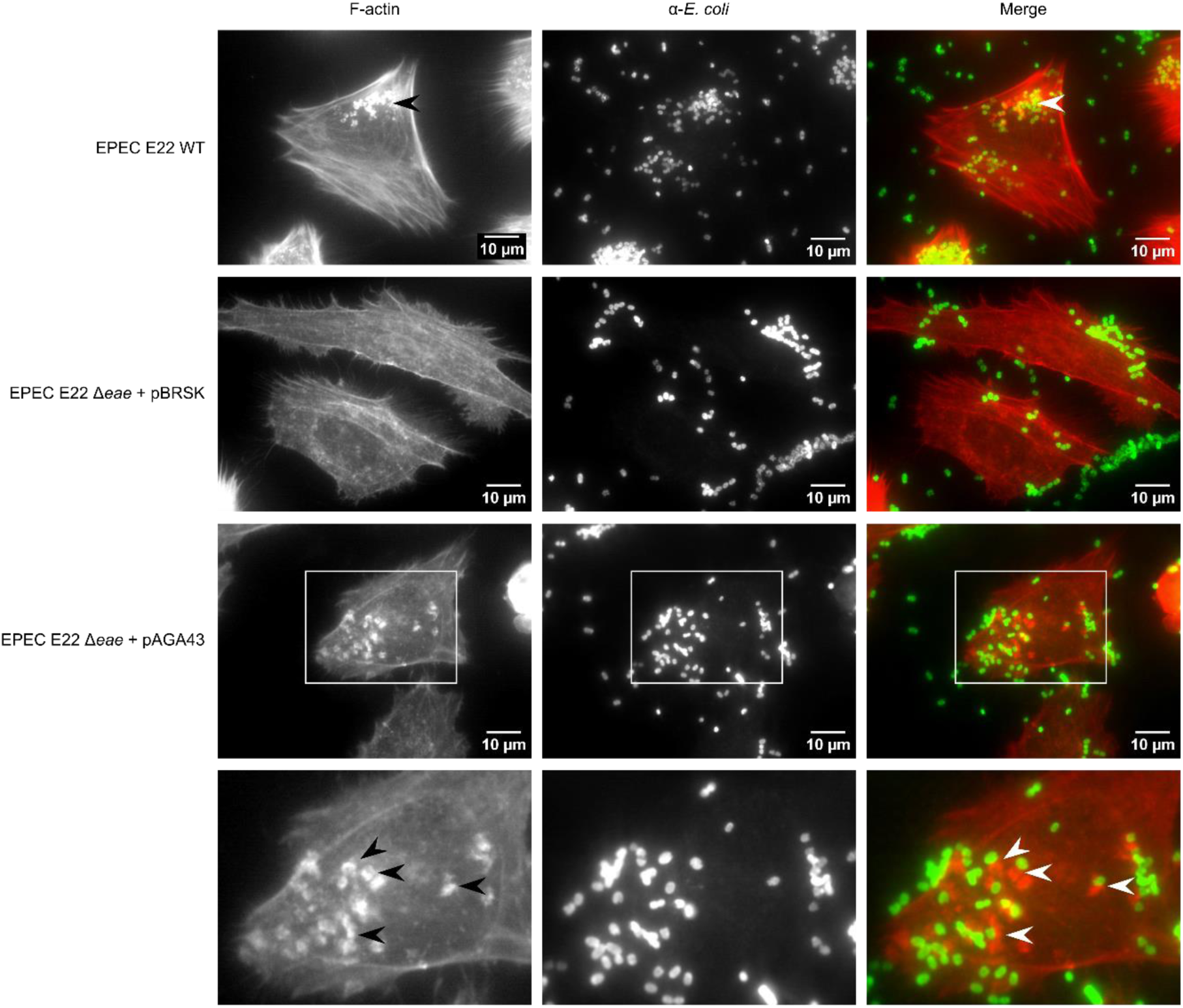
Int280 fused to Lpp-OmpA protein is addressed to the bacterial surface and restores the actin polymerization phenotype of the EPEC E22 intimin-mutant strain. Actin polymerization induced by the EPEC E22 strain in HeLa cells was observed with the Fluorescent Actin Staining (FAS) assay. HeLa cells were infected 5h with EPEC E22 WT, EPEC E22 Δ*eae* + pBRSK or EPEC E22 Δ*eae* + pBRSK_LO-Int280 (pAGA43) strains at a MOI of 50 bacteria per cell. Following staining of F-actin with rhodamine-phalloidin and bacteria with anti-*E. coli* antibodies, images were captured by epifluorescence microscopy. The lower panels show enlarged microscopy fields. Actin polymerization in some pedestals (red) beneath the bacteria (green) is indicated by arrowheads. Images are representative of three independent experiments.

### The OMV-Int280 vaccine elicits an IgG immune response in mice against the intimin of *C. rodentium* ICC169

To evaluate its potential, the OMV-Int280 vaccine was tested *in vivo* in mice to assess its innocuousness and efficacy at inducing an immune response against the intimin of *C. rodentium* ICC169. To increase the vaccine efficacy, a combination of OMVs containing luminal and surface-exposed Int280 at a ratio 1:1 (protein:protein) was used for vaccination. C57BL/6 female mice were vaccinated three times intraperitoneally (Fig. 3A), as this experimental procedure was previously shown to be efficient with mice vaccinated with purified Int280 (38) or OMVs enriched with antigens of interest (29, 39–41). A dose of mixed OMVs (0.5 µg of OMV- P-Int280 and 0.5 µg OMV-LO-Int280) or PBS (control) was used for each inoculation. A loss of 12.7 %, 9.9 % or 5.5 % of body weight was observed in vaccinated mice after the first immunization, the first and second booster, respectively, while mice in the PBS group did not lose weight (Fig. S1). Each weight loss was regained 2 to 3 days after each vaccination. No other clinical signs were observed in immunized mice. One mouse in the PBS group died unexpectedly (Fig. S1).

**FIG 3.**
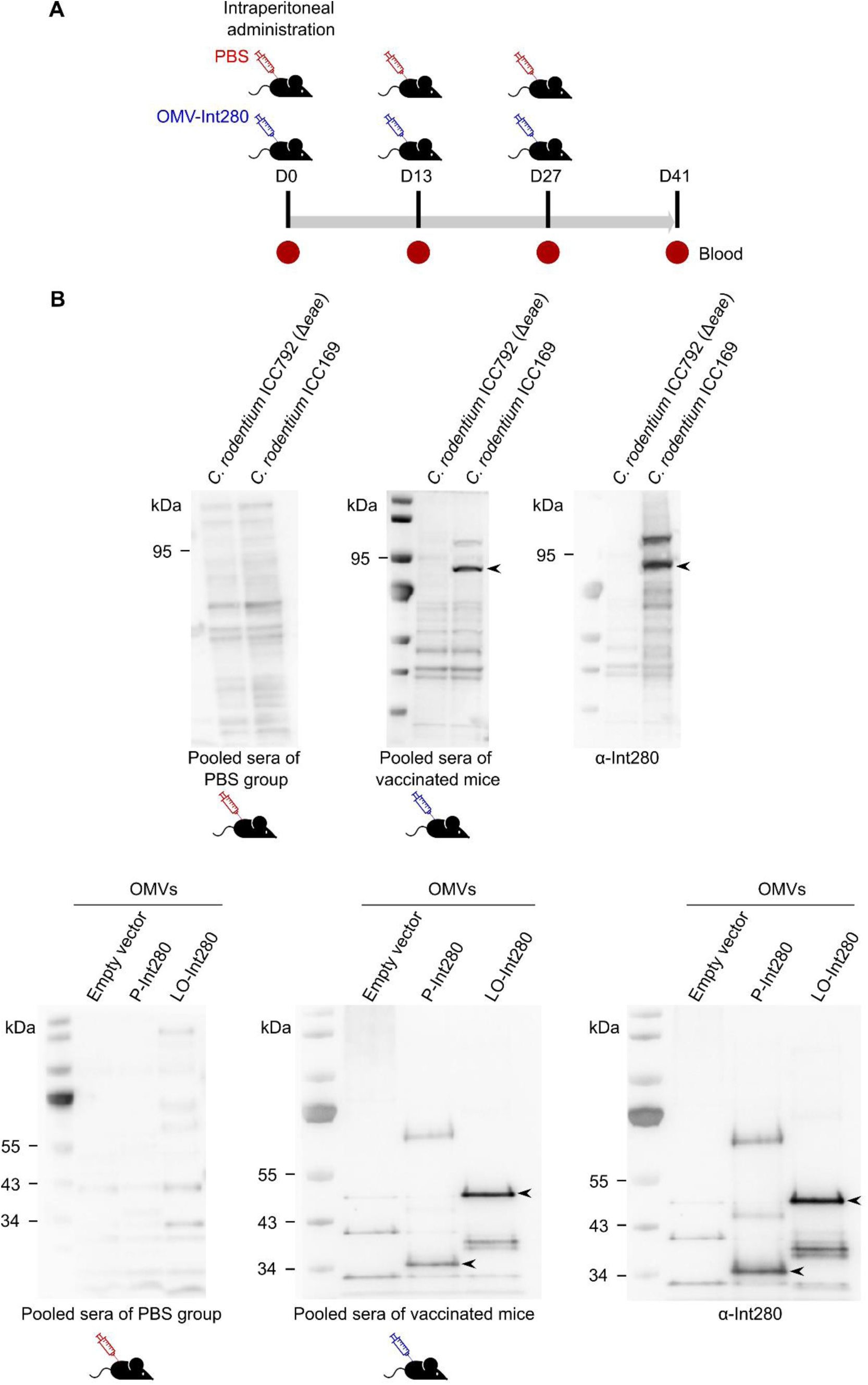
The OMV-Int280 vaccine elicited an anti-intimin IgG immune response in mice. (**A**) Experimental timeline of immunization experiment with female C57BL/6 mice. One group of 5 mice received PBS as a control and one group of 5 mice received the OMV-P-Int280/OMV-LO-Int280 mixed vaccine (OMV-Int280). The syringes indicate the days of immunization. Blood samples were collected on the indicated days for serum preparation. (**B**) Seroconversion of mice observed by Western Blotting with bacterial lysates of *C. rodentium* ICC792 (Δ*eae*) or *C. rodentium* ICC169, or with OMV-P-Int280 (2 µg) or OMV-LO-Int280 (2 µg) extracts obtained from *E. coli* BL21(DE3) Δ*ompF* carrying plasmid pAGA12 or pAGA16, respectively. “Empty vector” corresponds to the OMVs extracts obtained from *E. coli* BL21(DE3) Δ*ompF* carrying the empty pET-22b(+) plasmid. Sera collected from either PBS or vaccinated mice groups were each pooled and used to detect (i) full-length intimin (94kDa) from *C. rodentium* ICC169 bacterial lysates (top), or (ii) P-Int280 (30kDa) and LO-Int280 (43kDa) from OMVs fractions (bottom). The locations of intimin, P-Int280 and LO-Int280 are indicated with arrowheads. Anti-Int280 antibody was used as a positive control (top and bottom, third blot).

After the second booster, mice vaccinated with OMV-Int280 exhibited an IgG immune response against the full-length intimin of *C. rodentium* ICC169 and against both fusion proteins, i.e. P-Int280 and LO-Int280, contained in the vaccine (Fig. 3B). No such response was observed in mice that received PBS (Fig. 3B).

### The OMV-Int280 vaccine reduces the duration of fecal shedding of *C. rodentium* ICC169 in mice

Since the mixture of OMV-Int280 successfully induced a seroconversion against intimin, we further evaluated the effect of vaccination on the fecal shedding of *C. rodentium* ICC169 in mice.

Two weeks after the last booster, all mice from the immunized and control groups were orally challenged with 1x10^9^ CFU of *C. rodentium* ICC169. To monitor the shedding kinetics of *C. rodentium* ICC169, feces were collected at nine time points and subjected to *C. rodentium* enumeration (Fig. 4A). All mice shed *C. rodentium* ICC169 (Fig. 4B). Maximal *C. rodentium* load was reached between days 7 and 9 with approximately 1x10^8^ CFU/g of feces in all mice. On day 11, the mean *C. rodentium* shedding level of the immunized mice started to decrease, which was not the case for the control mice, whose shedding level started to decrease later, i.e. on day 14. In addition, the amount of *C. rodentium* in feces was significantly lower in the immunized group than in the PBS group from day 14 to day 18, suggesting intestinal clearance of *C. rodentium*. To verify the complete clearance of *C. rodentium* ICC169 in the vaccinated mice with a shedding level under the enumeration threshold, enrichment of their feces on day 14 and day 16 was performed and no *C. rodentium* was detected. On day 21, *C. rodentium* ICC169 shedding level was below the enumeration threshold in all but one mouse in both groups.

**FIG 4.**
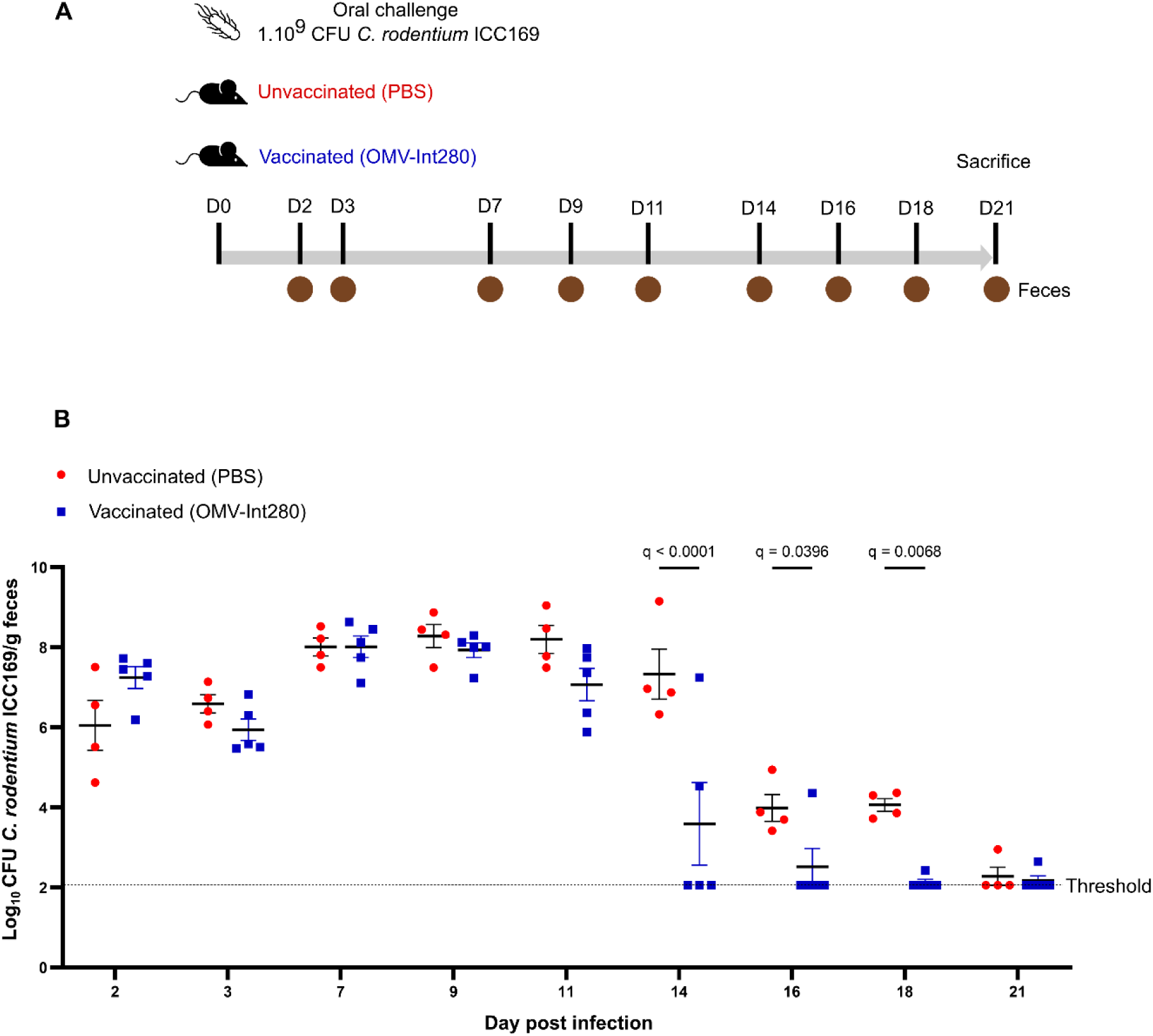
The OMV-Int280 vaccine reduced the duration of fecal shedding of *C. rodentium* in mice. (**A**) Experimental timeline of mice infection experiment. Feces were harvested at the indicated days. (**B**) Shedding kinetics of *C. rodentium* ICC169. Feces were weighted, homogenized, serially diluted and plated on Mac Conkey agar supplemented with nalidixic acid (50 µg.mL^-1^) to determine the *C. rodentium* load (log_10_ CFU/g of feces) for the PBS group (red circle) and the immunized group (blue square). The enumeration threshold was 2.05 log_10_ CFU/g feces. Statistical differences were considered significant when q-value < 0.05 (Two-way ANOVA).

## DISCUSSION

Two vaccines, Econiche® (Bioniche Life Sciences Inc.) (42) and Epitopix® (Vaxxinova) (43), have been developed and commercialized so far to reduce EHEC O157:H7 intestinal colonization in cattle. Econiche® vaccine is composed of Tir and effector proteins obtained from EHEC O157:H7 culture supernatants, while Epitopix® is based on siderophore receptor and porin (SRP) proteins purified from EHEC O157:H7 (42, 43). Experimental vaccines have also been developed targeting the intimin C-terminal domain of EHEC O157:H7 which is immunogenic (44–48) and triggers an immune response reducing EHEC fecal shedding in cattle (47, 48).

However, these commercialized or experimental vaccines are made from purified antigens or bacterial culture supernatants and require the addition of strong adjuvants to be efficient, rendering their production time-consuming and expensive (49).

The use of OMVs attracted increasing attention as a vaccinal platform due to their inability to replicate (50), spontaneous production by Gram-negative bacteria, immunogenic and adjuvant properties (21), reducing the cost and labor of vaccine production. In addition, OMVs are a versatile tool easy to load with antigens of interest (23–25). Here we exploited these advantageous properties for the development of an OMVs based vaccine containing Int280.

Two addressing systems derived from PelB and Lpp-OmpA proteins were used and shown to successfully export Int280 in the lumen and to the surface of OMVs, respectively. The hybrid protein Lpp-OmpA has previously been shown to successfully export the proteins monavidin (51) and SpyCatcher (52) to the surface of OMVs. Translocation into the lumen of OMVs of the Pfu DNA polymerase nanobody fused to the PelB signal sequence has also been described (53). In each case, those proteins were shown to be functional. Here, using the FAS assay, we demonstrated that the Lpp-OmpA system allowed the exposition of a functional Int280 protein to the bacterial OM surface.

Immunizations of C57BL/6 mice with the OMV-Int280 vaccine elicited a seroconversion against the intimin of *C. rodentium* ICC169. No clinical signs were observed after intraperitoneal administration of OMV-Int280, except for a body weight loss which might have been caused by the presence of LPS, since similar effects caused by LPS in mice have been described previously (54–56). In a previous work, subcutaneously immunization of mice with 10 µg of purified Int280 induced the production of anti-Int280 IgG1 and IgG2a. A production of IgG2a only was obtained from intranasal immunization (38). The oral and sublingual administration of C57BL/6 mice with a recombinant strain of *Lactobacillus casei* expressing a longer fragment of intimin (from AA363 to AA808, Int_CV_) (*L. casei*-Int_CV_) induced a mucosal immune response with the production of IgA (57).

This mouse model was also used here to determine whether vaccination could confer a reduction of fecal shedding of *C. rodentium* in mice C57BL/6. The shedding kinetics observed for *C. rodentium* ICC169 in the control PBS group was similar to those described in the literature (35, 58). Interestingly, the OMV-Int280 vaccine had an effect only on the recovery of mice during infection, with an earlier clearance of *C. rodentium* ICC169 observed in vaccinated mice compared to the control PBS group. An effect of intimin vaccines on mice colonization by *C. rodentium* has also been reported previously in C3H/HeJ mice vaccinated subcutaneously with purified Int280 (38) and in C57BL/6 mice immunized orally with *L. casei*-Int_CV_ (57). Our results thus confirm the interest of Int280 as an antigen to induce an immune response with anti-shedding properties.

Our study is a proof of concept intended to evaluate the OMV-Int280 vaccine safety and efficacy in a mice model. By showing elicitation of an immune response and a decrease of *C. rodentium* shedding duration in mice, this work brings a strong basis to further evaluate this vaccine in ruminants. However, as the digestive tract, immune system and response to endotoxin of ruminants are different from mice, the innocuousness, doses (number and concentration) and routes of administration of this OMV-Int280 vaccine will need to be assessed in ruminants.

Previous work has shown the lack of cross-protection between intimin variants carried by distinct EHEC serotypes as their C-terminal domains are antigenically different (38, 59), rendering crucial the use of specific intimin-variants to target peculiar

EHEC serotypes. In Europe, the EHEC serotype O26:H11 is the most common serotype isolated from patients with HUS (4). As this serotype possesses the β-intimin variant, the use of OMVs enriched with β-Int280 will be required to target the O26:H11 EHEC serotype in ruminants.

As a conclusion, OMVs were successfully loaded with luminal and surface Int280, providing a new type of vaccine with immunogenic properties leading to reduced fecal shedding of the targeted pathogen. This new type of vaccine based on OMVs and targeting Int280 is a promising candidate to reduce the intestinal carriage and fecal shedding of EHEC in ruminants and thus prevent foodborne EHEC infections.

## MATERIALS AND METHODS

### Bacterial growth conditions and plasmids

Bacterial strains and plasmids are listed in Table S1. All strains were precultured in Lennox L Broth Base (LB, Invitrogen, 12780029) overnight at 37°C, 240 RPM, with appropriate antibiotics (Carbenicillin [Euromedex, 1039-B], Nalidixic acid [Sigma-Aldrich, N4382] and Kanamycin [Euromedex, UK0015], each at 50 µg.mL^-1^).

The *E. coli* BL21(DE3) *ΔompF*::FRT mutant was constructed using the lambda Red recombination system, by first deleting a part of the *ompF* gene (from 47 to 945 bp) which was replaced by a kanamycin resistance gene amplified from pKD4 plasmid with the primers ompF-P1-M1 (5’-tagcaggtactgcaaacgctgcagaaatctataacaaagatggcaacaaaGTGTAGGCTGGAGCTGCT TC-3’) and ompF-P2-PM (5’-gttcaccagatcaacatcaccgataccttctacgtctttcgctttagattCATATGAATATCCTCCTTAG-3’).

Then, the kanamycin resistance cassette was removed using the pCP20 plasmid encoding the FLP recombinase (60).

### Construction of Int280 recombinant proteins

Plasmids used for the production of recombinant proteins were constructed by assembling DNA fragments amplified by PCR with primers listed in Table S2.

Except for the construction of plasmids pAGA12 and pAGA43 (see below), all the recombinant plasmids were constructed as follows. DNA amplicons were purified using the QIAquick® PCR Purification Kit (Qiagen, 28106) and assembled with pET-22b(+) DNA using the NEBuilder HiFi DNA Assembly Cloning Kit (New England Biolabs, E5520S) according to the manufacturer’s instructions. The resulting recombinant plasmids were introduced into NEB 5-alpha competent *E. coli* cells by thermal shock and transformed strains were incubated at 37°C for 1h. 100 µL were plated onto LB Agar containing carbenicillin (final concentration of 50 µg.mL^-1^) for selection and incubated at 37°C, overnight.

For the construction of plasmid pAGA12, the *eae*280 PCR product was purified using the QIAquick® PCR Purification Kit. It was digested with NcoI and BamHI restriction enzymes and ligated to NcoI/BamHI-digested pET-22b(+).The resulting plasmid was transformed into the TOP10 competent *E. coli* cells (C404010, Invitrogen) by thermal shock and transformed bacteria were selected as described above.

For the construction of plasmid pAGA43, the *eae* promoter (pEAE) was amplified from EPEC O103:H2 E22 genomic DNA with the primer pair AGA130-AGA131. The 1290 bp-long *lpp-ompA-eae280* fragment was amplified from pAGA16 with the primer pair AGA114-AGA132. The PCR fragments were purified using the QIAquick® PCR Purification Kit and amplified for producing a 1678 bp-long PCR fragment encoding the pEAE_*lpp*-*ompA*-*eae*280 fragment, containing the NotI and BamHI restriction sites at the 5’ and 3’end, respectively. The resulting PCR fragment was purified using the QIAquick® PCR Gel Extraction Kit (Qiagen, 28704) and cloned into the pCR XL-2- TOPO vector (Invitrogen, K8040-20) according to the manufacturer’s instructions. The resulting plasmid pCR XL-2-TOPO::pEAE_*lpp-ompA*-*eae*280 was introduced into OmniMAX competent *E. coli* cells (Invitrogen, C854003) by thermal shock and transformed strains were incubated at 37°C for 1h. 100 µL were plated onto LB Agar containing carbenicillin (final concentration of 50 µg.mL^-1^) for selection and incubated at 37°C, overnight. Following digestion of pCR XL-2-TOPO::pEAE_*lpp-ompA*-*eae*280 vector with NotI and BamHI restriction enzymes, the pEAE_*lpp-ompA*-*eae*280 fragment was purified and cloned into NotI/BamHI-digested pBRSK vector by ligation. The resulting recombinant plasmid was introduced into TOP10 competent *E. coli* cells by thermal shock and transformed strains were incubated at 37°C for 1h. 100 µL were plated onto LB Agar containing carbenicillin (final concentration of 50 µg.mL^-1^) for selection and incubated at 37°C, overnight.

Inserts were verified by Sanger sequencing (Eurofins Genomics). The recombinant plasmids were then transformed into the *E. coli* BL21(DE3) Δ*ompF* strain, except for the pBRSK and pAGA43 plasmids which were transformed into the EPEC O103:H2 E22 Δ*eae* strain.

### Production and purification of outer membrane vesicles

Bacterial strains were precultured overnight in 5 mL of LB supplemented with 50 µg.mL^-1^ of carbenicillin. Then, they were diluted at an optical density of OD_600nm_ = 0.1 in 50 mL of Terrific Broth (TB) medium (Sigma-Aldrich, T0918) supplemented with 50 µg.mL^-1^ of carbenicillin in 500 mL Erlenmeyer flasks with deflectors (Pyrex, ref 1142/10), and grown at 37°C with shaking at 240 RPM. After 2h of culture, gene expression was induced by the addition of isopropyl-β-D-thiogalactopyranoside (IPTG, Euromedex, EU0008-C) at a final concentration of 1 mM. After 4h of induction, bacteria were pelleted by centrifugation at 7,735 g for 10 min at 4°C. The supernatant was harvested and sterilized by filtration with 0.22 µm filter units (ClearLine, 257175). OMVs were then pelleted by ultracentrifugation at 175,000 g for 3h at 4°C, and resuspended in 50µL of phosphate buffered saline (PBS) supplemented with 0.1 g.L^-1^ of magnesium and calcium (Eurobio, CS1PBS00-01). For mice experiments, OMVs were resuspended in 50µL of PBS without magnesium and calcium (PBS) (Sigma-Aldrich, D8537). OMVs concentration was estimated by measuring the protein concentration with the BCA protein assay kit using bovine serum albumin (BSA) as a standard (DC™ Protein Assay Kit II, 5000112, Bio Rad).

### SDS-PAGE and Western blotting

Bacterial extracts of *C. rodentium* ICC169 (wild-type) and ICC792 (Δ*eae*) (Table S1) were prepared from cultures as follows: after 6 hours of growth in 20 mL of TB with appropriate antibiotics at 37°C and shaking at 240 RPM, 1 mL of bacterial cells at OD_600nm_ = 1 were collected and pelleted by centrifugation at 16,100 g for 2 min, washed with 1 mL of PBS, pelleted again, resuspended in 100 µL of loading buffer (NuPAGE Sample Reducing Agent [Invitrogen, NP0009] diluted at 1:10 in NuPAGE LDS Sample Buffer [Invitrogen, NP0007]) and sonicated. OMVs (2-10 µg) prepared from 50 mL cultures of *E. coli* BL21(DE3) Δ*ompF* as described above were diluted into a final volume of 10 µL of loading buffer. Samples were then boiled at 95°C for 5 min before being separated by SDS-PAGE (Invitrogen, NP0321) and either stained with Coomassie Blue (Sigma-Aldrich, B0770) or transferred to nitrocellulose membranes (Bio-Rad, 1704271) for Western blotting. Membranes were blocked with 3% BSA (Dominique Dutscher, P6154) in Tris Buffered Saline (TBS, Euromedex, ET220-B) with 0.1% of Tween® 20 (Sigma-Aldrich, P1379) (TBST) overnight at 4°C for Int280 and Intimin detection.

Membranes were probed with either (i) 1 µL of rabbit polyclonal anti-Int280β IgG (diluted 1:5000 in TBST-0.1% BSA) for 2h at RT, or (ii) 5 µL of mice sera of each group pooled, diluted in 5 mL in TBST-3% BSA and saturated with 200 µL of sonicated extracts of *C. rodentium* ICC792 (Δ*eae*) overnight at 4°C. Then, horseradish peroxidase (HRP)-conjugated goat anti-rabbit IgG antibody (diluted 1:5000 in TBST) (Bethyl Laboratories, A120-201P) or HRP-conjugated goat anti-mouse IgG antibody (diluted 1:5000 in TBST) (Bethyl Laboratories, A90-116P) were added for 1h at RT. Secondary antibodies were visualized using chemiluminescent peroxidase substrate detection reagents (Sigma-Aldrich, CPS3100) and imaged with a ChemiDoc XRS+ System (Bio-Rad).

### Fluorescent Actin Staining (FAS) assay

HeLa cells were seeded at 5.10^4^ cells per well in Lab-Tek 8 chambers slides (ThermoFisher Scientific, 154941) and cultivated 24h in Dulbecco’s Modified Eagle Medium (DMEM) with 4.5 g.L^-1^ of D- glucose, GlutaMAX^TM^ and 0.11 g.L^-1^ of sodium pyruvate (Gibco, 10569010) supplemented with 10 % of fetal calf serum (FCS) (Eurobio, CVFSVF00-01) and 1 % of non-essential amino acids (Sigma-Aldrich, M7145).

Bacterial strains EPEC O103:H2 E22, EPEC O103:H2 E22 Δ*eae* + pBRSK and EPEC O103:H2 E22 Δ*eae* + pBRSK_LO-Int280 were cultured in the activation medium (DMEM with 1 g.L^-1^ of D-glucose and 0.11 g.L^-1^ of sodium pyruvate [Gibco, 31966- 021], supplemented with 5 % of FCS, 1 % of D-(+)-mannose [Sigma-Aldrich, M6020], 25 mM of HEPES [Gibco, 15630080]) and IPTG at a final concentration of 1 mM for 4h at 37°C, with agitation at 240 RPM.

Prior to infection, HeLa cells were washed once with PBS and 500 µL of activation medium were added per well. Then, cells were infected with the bacteria at a multiplicity of infection (MOI) of 50 and centrifuged at 300 g for 5 min. After 3h of infection, the medium was replaced with 500µL of fresh activation medium per well and the infection continued for 2h. Cells were washed twice with PBS and fixed with 4 % formaldehyde (Electron Microscopy Sciences, 15713) in PBS for 12 min at RT. Cells were washed twice with PBS, permeabilized with 100 µL of Maxblock (Active Motif, 15252) supplemented with 0.3 % of Triton (Sigma-Aldrich, X100), and washed twice again with PBS. Bacteria were probed with 5 µL of rabbit anti-*E. coli* IgG antibody (Abcam, AB137967) diluted 1:200 in Maxblock and 100 µL were added per well, for 1h30 at RT. Then, cells were washed twice with 500µL of PBS supplemented with 0.05 % of Triton. The solution of secondary antibody was prepared by supplementing 1 mL of Maxblock with 1 µL of Alexa Fluor 488 goat anti-rabbit IgG antibody (diluted 1:1000) (Invitrogen, A11034) and 2 µL of rhodamine-phalloidin 400X (diluted 1:500) (Invitrogen, R415) to label EPEC and F-actin, respectively. 100 µL of this solution were added per well for 1h30, at RT and protected from light. Cells were washed twice with PBS supplemented with 0.05 % of Triton. Then, wells were removed and a coverslip was added on the slide with Fluoroshield (Sigma-Aldrich, F6182). The cells were observed using an Olympus BX41 microscope equipped with a 100x objective and a Leica DCF300FX camera. Epifluorescence images were acquired using a green filter (excitation 488 nm and emission 510 nm) and a red filter (excitation 540 nm and emission 565 nm) to observe EPEC and F-actin, respectively.

### Immunization of mice with OMVs-based vaccine and challenge with *Citrobacter rodentium* ICC169

The experimental protocol 2022092813461669 was approved by the Ethics Committee of the French Ministry of National Education and Higher Education. 4-week-old C57BL/6J female mice were purchased from Janvier Labs (Le Genest Saint Isle, France). C57BL/6 mice are considered resistant to *C. rodentium* infection with the development of a mild self-limiting disease with low inflammation and intestinal pathology(61, 62). All mice were housed in ventilated cages (five mice per cage) with a 12-hours day/night cycle, 20–24 °C temperature, 30–50% humidity with food and water *ad libitum*. Before the experiment, the mice were acclimated during two weeks.

In accordance with the European Directive 2010/63/EU on the protection of animals used for scientific purposes, the procedures performed for the study were minimally painful for the mice. However, for their comfort, sedation prior to blood collection and intraperitoneal administrations were achieved by inhalation with 4 % isoflurane at 2 L air/min.

After being anaesthetized by isoflurane inhalation, five 6-week-old mice were immunized three times intraperitoneally, at two-week intervals with 1 µg of a mixture of OMV-P-Int280 (0.5 µg) and OMV-LO-Int280 (0.5 µg) in a final volume of 100 µL of PBS. Five mice received 100µL of PBS in the control group. We vaccinated *via* the intraperitoneal route as this route of OMVs administration has been described to efficiently protect mice against pathogens (29, 39, 41). Blood samples were collected once a week by facial venipuncture under isoflurane inhalational anesthesia. Blood samples were centrifuged at 6,000 g for 10 min and the supernatants containing the sera were collected, aliquoted and stored at −20°C until use.

Two weeks after the last booster, all mice were challenged orally once, with 100µL of PBS containing 1x10^9^ CFU of *C. rodentium* ICC169. The inoculum of *C. rodentium* ICC169 was prepared as follows. *C. rodentium* ICC169 strain was grown overnight in LB supplemented with 50 µg.mL^-1^ of nalidixic acid, diluted (1:50) in 40 mL of LB supplemented with 50 µg.mL^-1^ of nalidixic acid and further grown 4 hours at 37°C with shaking at 240 RPM. Bacteria were then pelleted by centrifugation at 4,951 g for 10 min at RT and resuspended in 40mL PBS. OD_600nm_ was measured to calculate the volume of PBS required to resuspend *C. rodentium* ICC169 at a final concentration of 1.10^10^ CFU/mL. Then, bacteria were centrifuged again at 4,951 g for 10 min at RT and resuspended in the adequate volume of PBS. The inoculum of *C. rodentium* ICC169 was verified by retrospective plating on LB agar containing nalidixic acid (50 µg.mL^-1^). Shedding kinetics of *C. rodentium* ICC169 was monitored by plating feces onto Mac Conkey Agar supplemented with nalidixic acid (50 µg.mL^-1^), as follows. Feces were harvested before infection to verify that mice were not infected and then at multiple days after infection, in Lysing Matrix D tubes (MPBio, 116913050-CF) containing 500 µL of PBS. They were weighted and homogenized prior to serial dilution in PBS and plating. *C. rodentium* colonies were enumerated to determine the bacterial load in log_10_ CFU of *C. rodentium* ICC169/grams (g) of feces.

Mice were euthanized by cervical dislocation in agreement with the European directive 2010/63/EU on the protection of animals used for scientific purposes.

### Statistical analyses

The enumeration results are presented as the mean ± standard error of the mean (SEM). Statistical analysis was performed using GraphPad Software (La Jolla, CA, USA). Differences between groups were analyzed by two-way ANOVA with Benjamini, Krieger and Yekutieli comparison test. P-values were false discovery rate (FDR) corrected and a q-value < 0.05 was considered statistically significant.

## SUPPLEMENTAL MATERIAL

### Supplementary information

Table S1, Table S2 and Figure S1

## DATA AVAILABILITY

All data supporting the findings of this study are available within the article and its supplementary information file.

## ACKNOWLEDGEMENTS

We thank Gad Frankel (Imperial College, London) for the gift of the *C. rodentium* ICC169 and ICC792 strains and the Int280β antibody, George Georgiou (Department of Chemical Engineering, University of Texas, Austin) for the pTX101 vector and Mathieu Epardaud (INRAE, Université de Tours) for critical reading of the manuscript. This project was supported by a grant from the French National Research Agency (ANR) (PreventEHEC, ANR-21-CE21-0006-01_ACT) and by a doctoral grant from the French National Research Institute for Agriculture, Food and Environment (INRAE, France).

A.GA., C.G., C.S., P.M., A.GO., J-P.N., F.A. and P.B. conceptualized the study. A.GA., F.A. and P.B. analyzed the study data. A.GA. wrote the original draft of the manuscript. A.GA., J-P.N., E.O., F.A. and P.B. reviewed and edited the manuscript. F.A. and PB acquired funding for this study. E.O., F.A. and P.B. supervised the project

